# Wnt target gene Ascl4 is dispensable for skin appendage development

**DOI:** 10.1101/2024.01.12.575337

**Authors:** Verdiana Papagno, Ana-Marija Sulic, Jyoti P. Satta, Aida Kaffash Hoshiar, Vinod Kumar, Jukka Jernvall, Marja L. Mikkola

## Abstract

The development of skin appendages, including hair follicles, teeth and mammary glands is initiated through the formation of the placode – a local thickening of the epithelium. The Wnt/β-catenin signaling cascade is an evolutionary conserved pathway with an essential role in placode morphogenesis, but its downstream targets and their exact functions remain ill defined. In this study, we identify *Achaete-scute complex-like 4* (*Ascl4*) as a novel target of the Wnt/β-catenin pathway and demonstrate its expression pattern in the signaling centers of developing hair follicles and teeth. Ascl transcription factors belong to the superfamily of basic helix-loop-helix transcriptional regulators involved in cell fate determination in many tissues. However, their specific role in the developing skin remains largely unknown. We report that *Ascl4* null mice have no overt phenotype. Absence of Ascl4 did not impair hair follicle morphogenesis or hair shaft formation suggesting that it is non-essential for hair follicle development. No tooth or mammary gland abnormalities were detected either. We suggest that other transcription factors may functionally compensate for the absence of Ascl4, but further research is warranted to assess this possibility.

## Introduction

The development of ectodermal appendages, such as hair follicles, teeth and various glands, shares a set of well-defined stages, including induction, placode and bud stages (Mikkola & Millar, 2006). In the early stages of development, reciprocal and sequential tissue interactions between the ectodermal epithelium and the underlying mesenchyme result in the formation of an epithelial thickening, or placode, serving as a precursor for the mature ectodermal appendage (Biggs & Mikkola, 2014; Mikkola & Millar, 2006). Once placodes are established, the epithelium expands into the condensing mesenchyme to form a bud, followed by organ-specific epithelial morphogenesis. The specific details of the later morphogenetic and differentiation programs vary widely among different organs, reflecting the diverse forms and functions of fully formed ectodermal appendages (Biggs & Mikkola, 2014). The invaginating hair bud progresses via peg and bulbous peg stages, concomitant with gradual differentiation of the follicular cell types, to produce the mature hair follicle that undergoes cyclic regeneration throughout the lifetime (Sennett & Rendl, 2012). Tooth morphogenesis, on the other hand, proceeds through the cap and bell stages, prior to the onset of mineralization of the hard tissues and root formation (Balic & Thesleff, 2015). The mammary bud initially grows slowly, but when it reaches the adjacent fat pad precursor tissue (prospective adult stroma) it undergoes branching morphogenesis generating a small ductal tree by birth (Spina & Cowin, 2021).

Ectodermal appendage development is orchestrated by a limited number of conserved signaling cascades, encompassing both shared and unique pathways for each organ (Mikkola & Millar, 2006). Members of the Wnt, transforming growth factor β (Tgfβ), fibroblast growth factor (Fgf), hedgehog (Hh), and tumor necrosis factor (Tnf) families have been extensively studied during ectodermal organ formation using genetically engineered mouse models. However, the downstream events and target genes remain incompletely characterized (Biggs & Mikkola, 2014; Yu & Klein, 2020). The canonical Wnt pathway, mediated by β-catenin and Tcf/Lef1 family of transcription factors, is an evolutionary conserved pathway with essential roles in tissue morphogenesis and cell fate specification (Loh et al., 2016). In developing skin appendages, it plays a crucial role in the early inductive events. The suppression of canonical Wnt signaling in the epidermis, achieved either by overexpression of the secreted Wnt inhibitor Dkk1 or epithelial deletion of β-catenin, leads to absence of all signs of hair and mammary placode formation (Andl et al., 2002; Chu et al., 2004; Huelsken et al., 2001; Zhang et al., 2009), and tooth development halts at the placode stage (Liu et al., 2008). On the other hand, forced epithelial β-catenin activation in the epidermis leads to precocious induction of hair follicle development, committing the entire skin to hair follicle fate (Närhi et al., 2008; Zhang et al., 2008), and in the oral epithelium, results in the formation of supernumerary teeth (Järvinen et al., 2006; Liu et al., 2008).

The acquisition of specific morphogenetic programs not only requires essential signaling molecules but also distinct combinations of transcription factors that elicit cell fate changes during embryonic development (Iwafuchi-Doi & Zaret, 2016). *Ascl4* belongs to the achaete-scute complex-like family of transcription factors (*Ascl*1-5), characterized by a conserved basic-helix-loop-helix DNA-binding domain (García-Bellido & De Celis, 2009). Various *Ascl* family members are involved in cell fate determination in different tissues such as *Ascl1* in neural fate commitment (Guillemot & Hassan, 2017), and *Ascl2* in intestinal stem cell and trophoblast cell lineage specification (Lefebvre, 2012; van der Flier et al., 2009). However, knowledge about *Ascl4* is limited, although it is known to be expressed in the embryonic human skin (Jonsson et al., 2004).

Recently, we transcriptionally profiled mouse skin epithelial cells at embryonic day 14.5 (E14.5) when the first hair placodes have just formed. Our bulk RNA sequencing (RNA-seq) data, confirmed by unbiased single-cell RNA-seq (scRNA-seq) profiling, revealed that *Ascl4* is highly enriched in the hair placode population compared to the interfollicular epidermis (Sulic et al., 2023), consistent with findings from other hair placode profiling studies (Sennett et al., 2015; Tomann et al., 2016). In this study, we investigated the role of Ascl4 in ectodermal appendage development, specifically focusing on hair follicles, teeth, and mammary gland. Our findings establish a connection between Ascl4 and Wnt signaling, showing that *Ascl4* is a likely direct transcriptional target of the Wnt pathway. However, our results indicate that Ascl4 is dispensable for ectodermal organogenesis.

## Results

### *Ascl4* is expressed in hair and tooth signaling centers

Our recent scRNA-seq profiling of hair placode-enriched cells identified four spatially distinct cell populations: one representing a central population (Placode-I), another one at the placode edge (Placode-IV), and two populations located in-between (Placode-II and Placode-III). Intriguingly, *Ascl4* was predominantly expressed in the Placode-I population (Fig. 1A; Sulic et al., 2023). This cell population is defined by high expression of signaling molecules such as *Tgfβ2, Bmp2*, and *Shh*. Other markers include *Tgfb2* and *Lhx2*, as well as *Sp5* (Fig. 1A), a direct target gene of the Wnt/β-catenin pathway (Takahashi et al., 2005; Weidinger et al., 2005). Interestingly, *Axin2*, commonly used as the readout of the canonical Wnt signaling activity, was also expressed at high levels in Placode-I (Fig. 1A).

**Figure 1.**
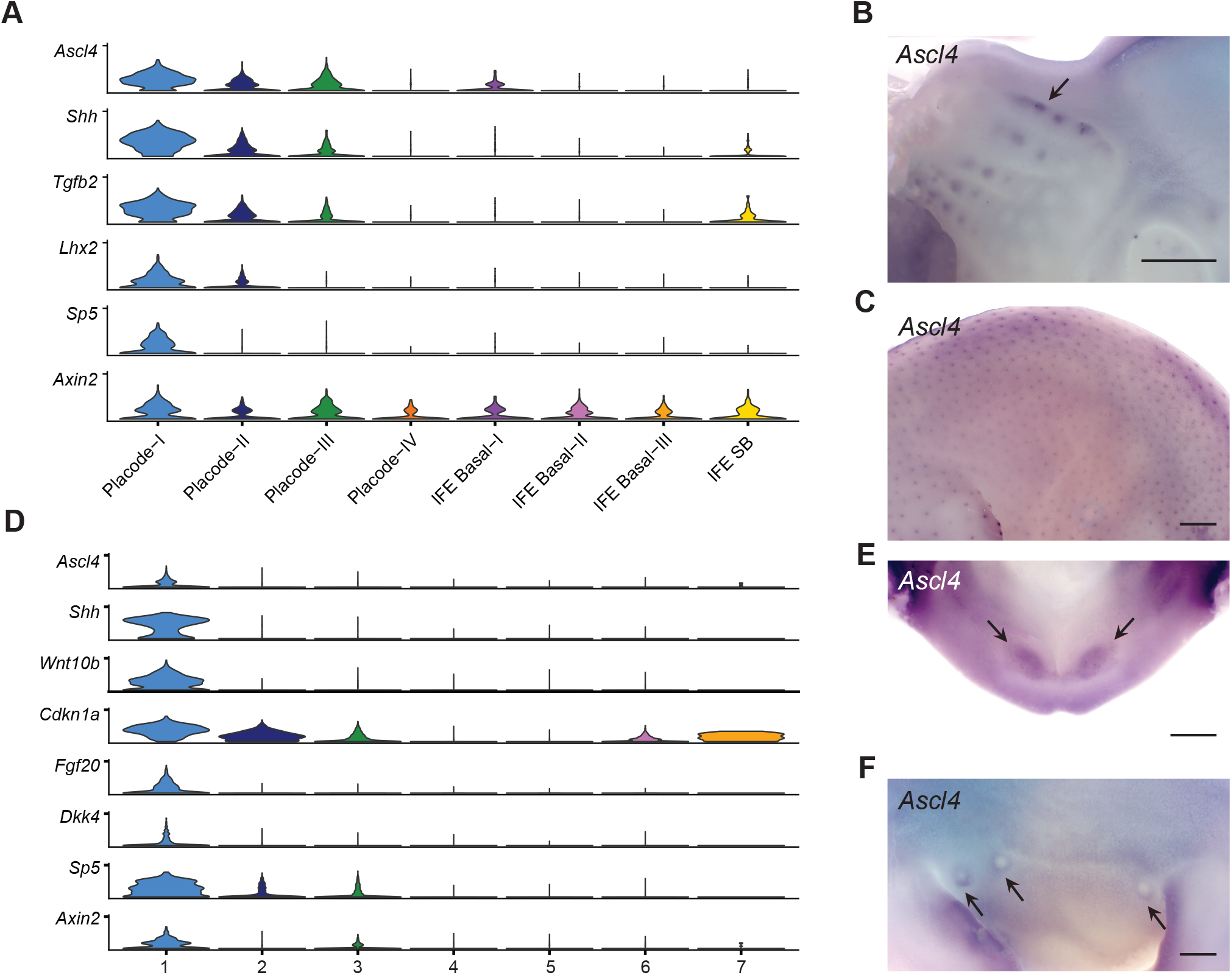
*Ascl4* is expressed in signaling centres of hair follicle and teeth. **(A)** Stacked violin plots showing expression of selected genes in E14.5 hair placode-enriched scRNA-seq dataset (Sulic et al., 2023). Placode-I to Placode-IV clusters represent genuine placode populations, while other clusters represent various interfollicular cell populations, as detailed in Sulic et al., 2023. **(B-C)** Expression of *Ascl4* was analyzed by whole mount *in situ* hybridization in E12.5 heads (n=5) **(B)** and E14.5 embryos (n=8) **(C)**. Arrow in B indicates expression of *Ascl4* in the whisker placodes. Scale bar, 500 μm. **(D)** Stacked violin plots showing expression of *Ascl4*, and selected enamel knot markers in the epithelial cell populations of E14.5 molar tooth scRNAseq dataset (Hallikas et al., 2021). **(E-F)** Expression of *Ascl4* was analyzed by whole mount *in situ* hybridization in E12.5 lower jaw (n=7) **(E)** and E12.5 mammary buds (n=8) **(F)**. Scale bar, 200 μm. Arrows indicate *Ascl4* expression in incisor and mammary buds, respectively.

To analyze *Ascl4* expression during the induction of hair and vibrissae development *in situ*, we performed whole-mount *in situ* hybridization (WMISH). At E12.5, *Ascl4* expression was detected in the whisker placodes (Fig. 1B). By E14.5, *Ascl4* mRNA was also localized in the hair placodes of back skin (Fig. 1C) (Sulic et al., 2023). At both developmental stages, *Ascl4* expression appeared to be restricted to epithelial compartment, consistent with transcriptomic profiling of E14.5 whole skin by bulk and scRNA-seq (hair-gel.net, Sennett et al., 2015; kasperlab.org/embryonicskin, Jacob et al., 2023).

Many genes and pathways have shared functions and similar expression patterns across various ectodermal appendages (Mikkola & Millar, 2006), prompting us to explore *Ascl4* expression also in the embryonic tooth and mammary gland. We have recently generated scRNA-seq data on developing molar teeth at E14.5 (Hallikas et al., 2021), a stage characterized by the appearance of the enamel knot, a signaling center governing tooth cusp patterning (Jernvall & Thesleff, 2000). Analysis of *Ascl4* expression in this dataset revealed that it was restricted to the epithelium, in a rather small cluster of cells (Fig. S1A). Further sub-clustering of the epithelial cells highlighted *Ascl4* enrichment in a cell population defined by high expression of *Shh, Wnt10b, Cdkn1a, Fgf20, Dkk4*, and *Sp5* (Fig. 1D), all known markers of the enamel knot (Vaahtokari et al., 1996; Jernvall et al., 1998; Fliniaux et al., 2008; Porntaveetus et al., 2011; Ahtiainen et al., 2016). This identifies *Ascl4* as a novel enamel knot marker. Also *Axin2* was enriched in the same cell population (Fig. 1D), consistent with previous studies reporting high Wnt signaling activity in the enamel knot (Liu et al., 2008; Lohi, Tucker and Sharpe, 2010; Ahtiainen et al., 2016). To assess if *Ascl4* is expressed already during the tooth placode stage, we performed WMISH at E12.5. At this stage, *Ascl4* transcripts were detected in the incisor primordia, albeit at low levels (Fig. 1E). Finally, we explored the expression of *Ascl4* in the mammary bud. WMISH revealed weak *Ascl4* expression in the mammary bud area at E12.5 (Fig. 1F), specifically in a region surrounding the mammary buds, forming a ring-like pattern.

### *Ascl4* is a likely transcriptional target of the Wnt/β-catenin pathway

Since *Ascl4* expression was detected at sites of high Wnt activity, we next investigated whether it acts downstream of the Wnt pathway. To explore the link between *Ascl4* and Wnt signaling *in vivo*, we analyzed *Ascl4* expression in mice with forced epithelial activation of β-catenin (Närhi et al., 2008), expressing one wild-type and one exon 3 deleted allele of *Ctnnb1* (encoding a stabilized form of β-catenin) in the mutant skin epithelium. At E12.5, *Ascl4* transcripts were detected in the mammary line, as well as in ectopic hair placodes that begin to emerge at this stage (Fig. 2A). A day later, *Ascl4* was still undetectable in the back skin of control embryos, whereas its expression had intensified in the ectopic hair placodes and in mammary buds of stab-β-cat embryos. The latter finding is in line with our recent RNA sequencing data that revealed a 7.6-fold upregulation of *Ascl4* expression in stab-β-cat mammary buds compared to wild type at E13.5 (Satta et al., 2023).

**Figure 2.**
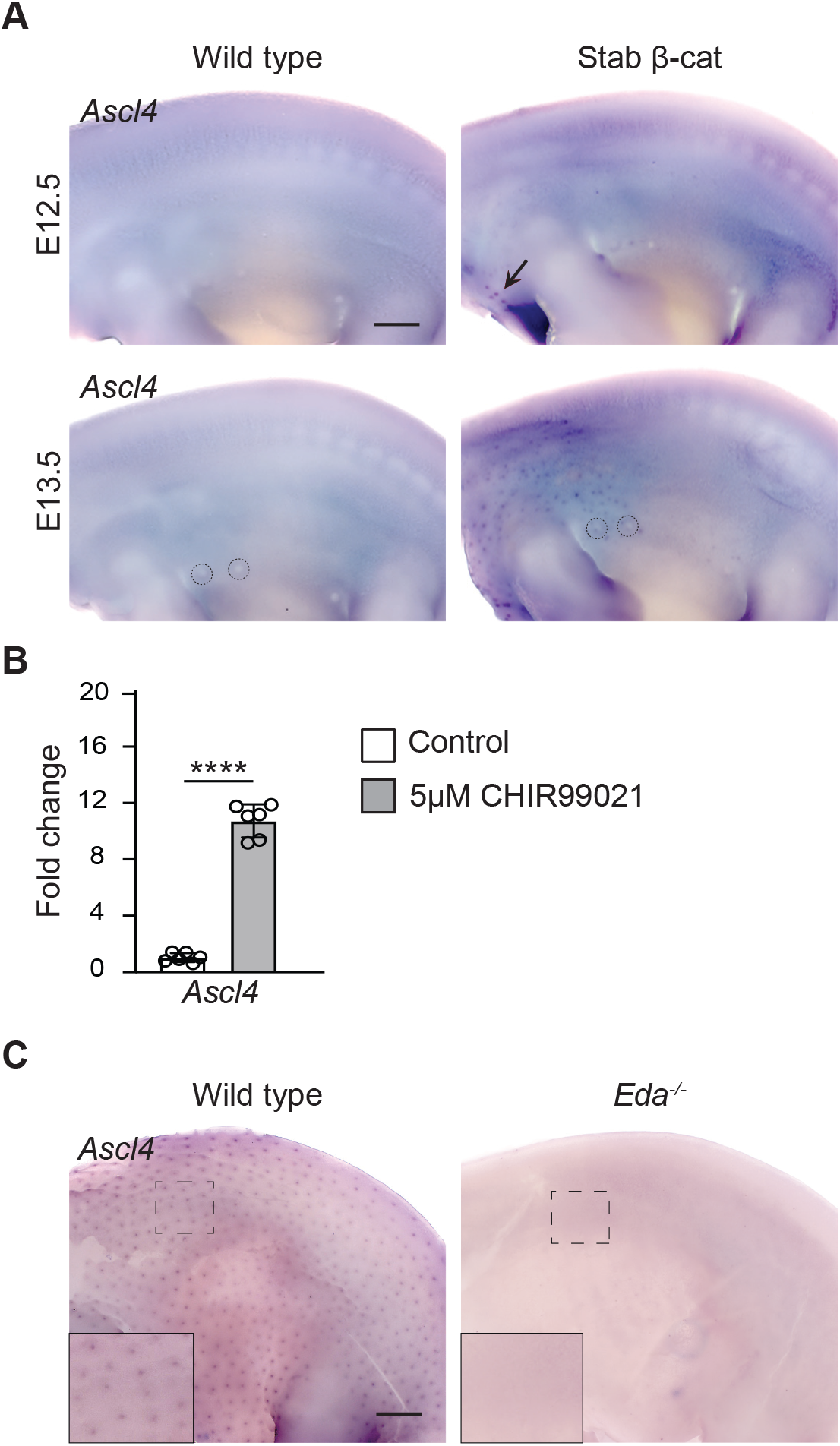
*Ascl4* is a Wnt target gene. **(A)** *Ascl4* expression in wild-type (n=3) and stabilized β-catenin (n=3) skin analyzed by whole mount *in situ* hybridization at E12.5 (top panels) and E13.5 (bottom panels). Arrow indicates the first hair placode-like structures that have appeared at E12.5. Dashed circles highlight mammary bud 2 and 3. Scale bar, 500 μm. **(B)** qRT-PCR of *Ascl4* after 4 hours treatment with 5μM CHIR99021 or vehicle (n=6). Data are shown as mean ± s.d. ****p < 0.001. Statistical significance was assessed with two-tailed unpaired Student’s t-test. **(C)** *Ascl4* expression in wild-type (n=4) and *Eda*^*-/-*^ embryos (n=4). Insets show higher magnification of the back skin. Scale bar, 500 μm.

These data prompted us to ask whether *Ascl4* could be a direct Wnt/β-catenin target gene. To this end, we investigated the ability of CHIR99021, a potent activator of the canonical Wnt pathway (Naujok et al., 2014), to induce *Ascl4* expression in E14.5 embryonic back skin explants. Explants were divided into two halves, one treated with vehicle and the other with 5μM CHIR99021, followed by expression analysis by qRT-PCR. 4-hr exposure to CHIR99021 led to a highly significant, 10.7-fold increase in *Ascl4* mRNA levels compared to controls (Fig. 2B).

The ectodysplasin (*Eda*)-Eda receptor (*Edar*) pathway lies downstream of Wnt signaling in hair placodes (Biggs & Mikkola, 2014). *Eda*-deficient primary hair placodes are characterized by low Wnt/β-catenin signaling activity and failure to proceed beyond a pre-placode stage (Schmidt-Ullrich et al., 2006; Fliniaux et al., 2008; Zhang et al., 2009). Some of the known Wnt target genes, such as *Dkk4* and *Fgf20*, are expressed in *Eda* null pre-placodes at E14.5, though at greatly reduced levels (Fliniaux et al., 2008; Huh et al., 2013), suggesting that these genes respond to even low Wnt/β-catenin activity, while many other placode markers such as *Shh*, and the Wnt target gene *Sp5* are undetectable (Huh et al., 2013; Sulic et al., 2023). Similarly, no *Ascl4* transcripts could be detected in the back skin of *Eda* null embryos at E14.5 (Fig. 2C).

### Ascl4 is not required for hair follicle morphogenesis

To investigate the function of *Ascl4* in ectodermal appendages, we used mice lacking Ascl4 (hereafter referred to as *Ascl4* KO mice). *Ascl4* KO mice were viable and fertile, indistinguishable from their wild-type and heterozygous littermates in size, weight, and external appearance (Fig. S1B-C). Mutant embryos did not exhibit any gross phenotypic differences and were obtained at Mendelian ratios, indicating that the deletion of *Ascl4* did not result in embryonic lethality (Fig. S1D).

In mice, hair follicles are induced in three successive waves, resulting in the production of four different hair types: the guard (first wave), awls and auchene (second wave), and zigzag (third wave) hairs (Sennett & Rendl, 2012). To assess the functional importance of *Ascl4* in hair follicle development, we first examined the expression of three well-characterized hair placode markers – *Dkk4, Wnt10b* and *Shh* – in *Ascl4* KO embryos using WMISH. At E13.5, *Dkk4* expression was readily detectable, while *Shh* was just starting to emerge, but no difference was observed between the genotypes (Fig. 3A). At E14.5, all genes were expressed in the primary hair placodes of *Ascl4* KO embryos, showing spatial and temporal expression comparable to wild-type embryos (Fig. 3A; Fig. S1E). Histological analysis did not reveal any abnormalities in placode morphology either (Fig. 3B). Similar to controls, at E16.5, primary hair follicles in *Ascl4* KO embryos had advanced to the peg stage, and secondary hair placodes had formed. By E18.5, all three waves of hair follicles were visible in the mutant skin, with no apparent changes in their sizes, lengths, and distribution (Fig.3B).

**Figure 3.**
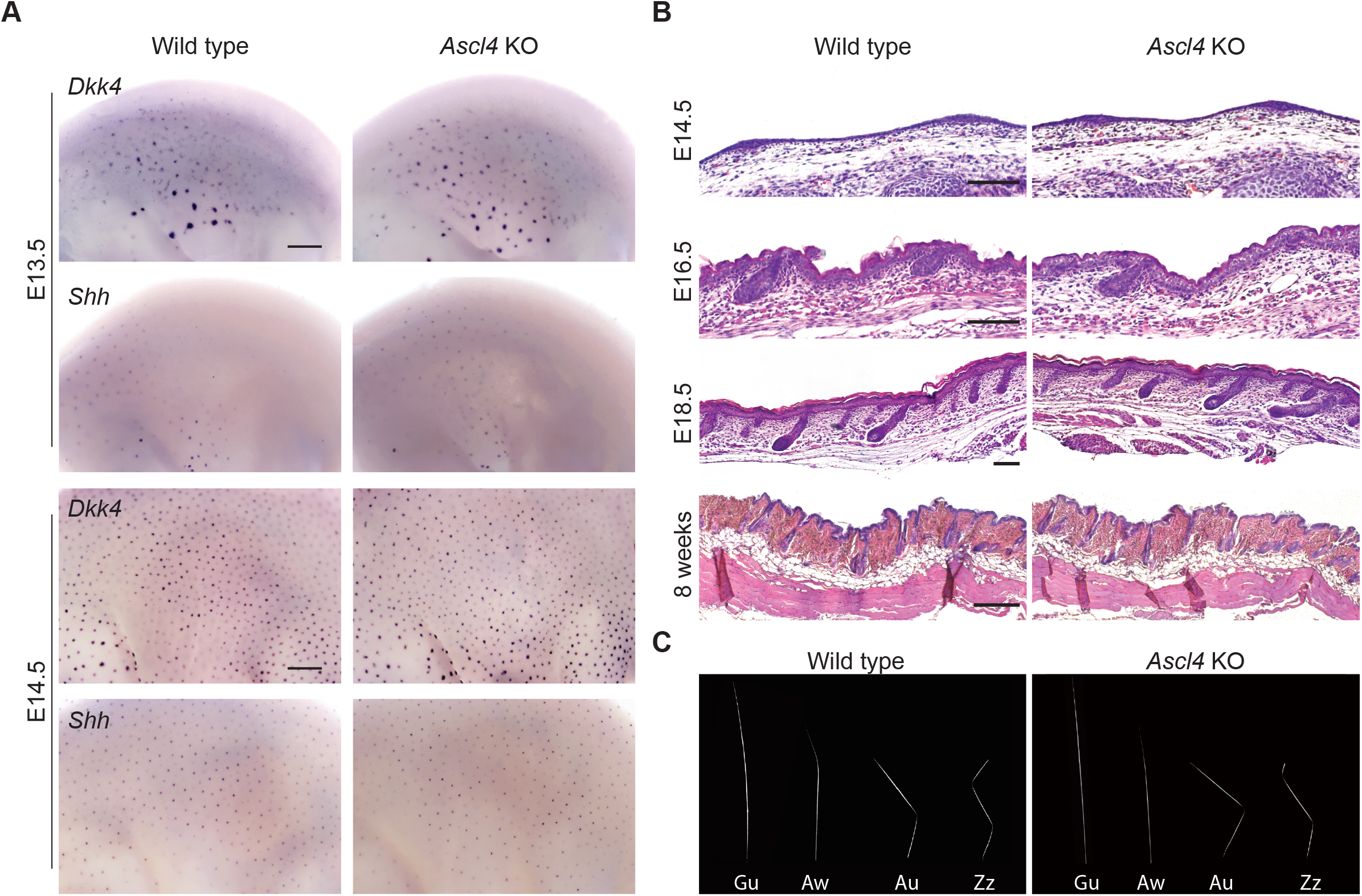
*Ascl4* is not required for hair follicle morphogenesis. **(A)** Expression of *Dkk4* and *Shh* was assessed by whole mount *in situ* hybridization in wild-type (n=4) and *Ascl4* KO (n=4) embryos at E13.5 (upper panel) and E14.5 (bottom panel). Scale bar, 500μm. **(B)** Hematoxylin and eosin staining of wild type and *Ascl4* KO E14.5, E16.5, E18.5, and 8-week-old dorsal skin (n=6 each). Scale bar, 100μm (E14.5, E16.5, E18.5), and 200μm (8-wk skin). **(C)** Hair types of wild type (n=6) and *Ascl4* KO mice (n=6) at 8 weeks of age. Gu, guard; Aw, awl; Au, auchene; Zz, zigzag.

Next, we analyzed whether loss of *Ascl4* would impact postnatal hair growth and progression of the hair cycle. Between postnatal day 15-25, hair follicles undergo their first catagen (destruction phase), and then a longer telogen (rest phase), commencing ∼ 7 weeks of age (Müller-Röver et al., 2001). No difference was observed in the hair follicles of wild-type and *Ascl4* KO mice at 8 weeks of age (Fig. 3B). The fur of *Ascl4* KO mice contained all four hair types, and the external hair structures were not affected either (Fig. 3C). Thus, the loss of *Ascl4* does not seem to impair hair follicle development, cycling, or hair shaft formation.

The lack of an obvious hair phenotype in *Ascl4* KO mice raised the question whether other *Ascl* genes could compensate for absence of *Ascl4*. Our bulk E14.5 RNA-seq data did not reveal the presence of *Ascl1* or *Ascl3* (sulic.rahtiapp.fi; Sulic et al., 2023), whereas *Ascl2* and *Ascl5* were detected in the hair placode population, albeit at much lower levels than *Ascl4* (Fig. S1F). However, *Ascl2* transcripts were enriched in the interfollicular epithelium, not the placode (Fig. S1F). These findings were confirmed by inspection of other published E14.5 databases (Sennett et al., 2015; Jacob et al., 2023). Also postnatally, *Ascl2* and *Ascl4* were expressed in non-overlapping areas, with *Ascl2* being enriched in the interfollicular epidermis, other *Ascl* genes being expressed at negligible levels in follicular cell populations (Skin-GLOW, https://rstudio-connect.hpc.mssm.edu/skinglow_minerva/).

### Ascl4 is not essential for mammary gland and tooth formation

Given that *Ascl4* was expressed also in developing teeth and mammary glands, we analyzed the tooth and mammary gland phenotype of *Ascl4* KO mice. The dentition of *Ascl4* KO mice appeared normal, with molar shapes and sizes indistinguishable from those of wild-type littermate controls (Fig. 4A). To investigate potential redundancy with other family members, we analyzed the expression of *Ascl* genes in the E14.5 molar tooth scRNA-seq dataset (Hallikas et al., 2021). Of the five family members, only *Ascl4* and *Ascl5* transcripts could be detected in a clustered manner, while other *Ascl* genes were expressed only in few sporadic cells (Fig. S1A, S1G). Interestingly, *Ascl4* and *Ascl5* displayed a similar expression pattern, showing enrichment in the enamel knot.

**Figure 4.**
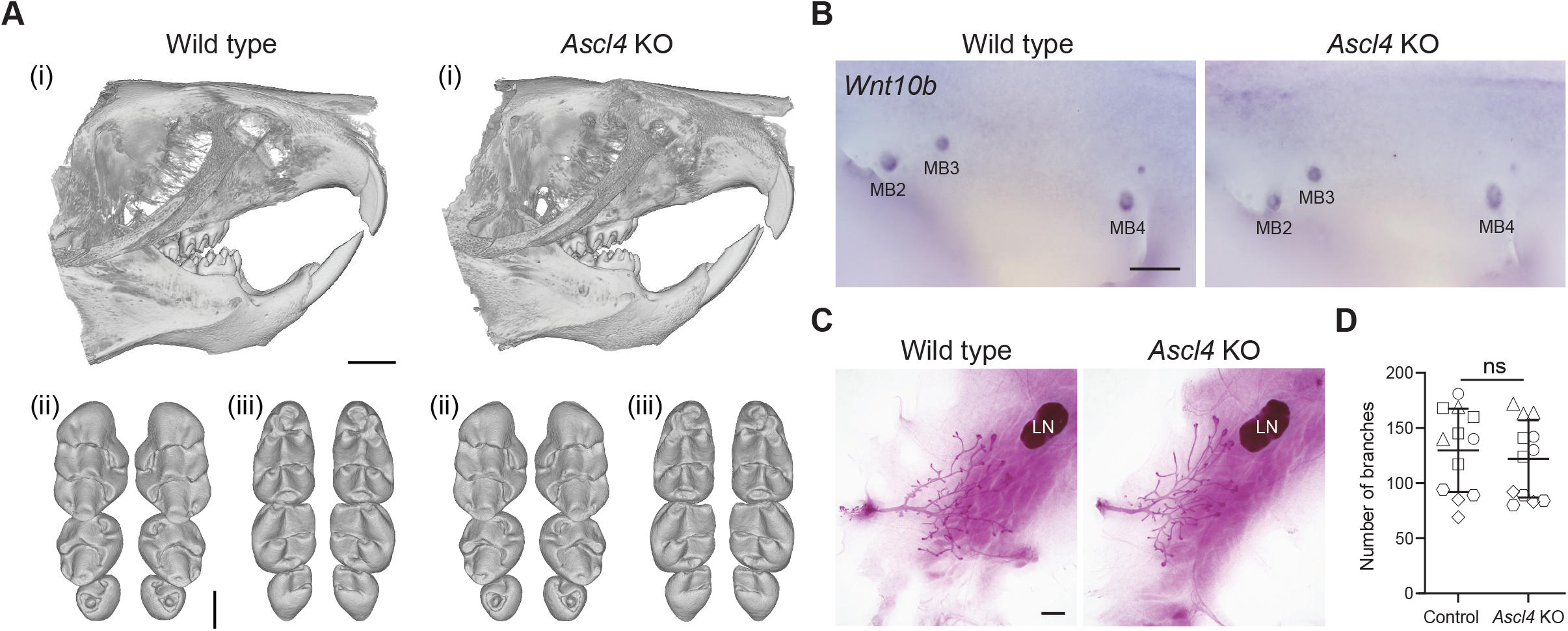
*Ascl4* is dispensable for mammary gland and tooth development. **(A)** Representative μCT scans showing incisor (i), and upper (ii) and lower molars (iii) of 4-week-old wild-type (n=6) and *Ascl4* KO mice (n=6). Scale bar, 1.5 mm (incisors), and 500 μm (molars). **(B)** Expression of the mammary bud marker *Wnt10b* in wild-type (n=5) and *Ascl4* KO (n=5) embryos at E12.5. MB, mammary bud. Scale bar, 500 μm. **(C)** Representative images of whole-mount carmine alum stained 4th mammary gland of 4-week-old wild-type and *Ascl4* KO mice. LN, lymph node. Scale bar, 1 mm. **(D)** Quantification of branches (ductal tips) of the 4th mammary gland of control (*Ascl4*^+/+^ and *Ascl4*^+/-^) (n=12) and *Ascl4* KO (n=12) mice at 4 weeks of age. The shape of the symbol indicates littermates. Statistical significance was assessed with two-tailed unpaired Student’s t-test. ns, not significant.

To characterize the *Ascl4* KO mammary phenotype, we first analyzed the expression of *Wnt10b*, a well-known mammary bud marker using WMISH, but found no obvious difference between wild-type and *Ascl4* KO embryos at E12.5 (Fig. 4B). The majority of mammary gland growth and branching takes place during puberty, which commences around 3 weeks of age (Macias & Hinck, 2012). The mammary ductal trees of 4-wk old *Ascl4* KO mice appeared morphologically indistinguishable from the wild-type littermates (Fig. 4C), a finding confirmed by quantifying the number of ductal tips (Fig. 4D). *Ascl4* KO females were also able to nurture their pups (n=6 females, 11 litters, 78 pups), suggesting the absence of any gross lactation defects.

## Discussion

The canonical Wnt/β-catenin pathway has long been known to be essential for ectodermal appendage development (Andl et al., 2002; Van Genderen et al., 1994). Here, we identify the transcription factor Ascl4 as a novel target of the Wnt/β-catenin pathway, co-expressed with *Shh* at signaling centers of developing teeth and hair follicles. Inspection of publicly available RNA-seq hair follicle datasets (accessed via Skin-GLOW, https://rstudio-connect.hpc.mssm.edu/skinglow_minerva/) indicates that also at later developmental stages, *Ascl4* is co-expressed with *Shh* in the transit amplifying cells (activated stem cells) of the hair follicle, as well as the hair matrix encompassing the proliferative precursors giving rise to the hair shaft.

Different levels of Wnt/β-catenin signaling often result in the activation of distinct target genes which in turn determine different cell fates in a context-dependent manner (Matos et al., 2020; Mourao et al., 2021; Söderholm & Cantù, 2021). Our data suggest that *Ascl4* expression may require high levels of Wnt/β-catenin activity. Unlike in hair follicles and molar teeth, *Ascl4* was not detected in the invaginating mammary bud but rather at its edges. This contrasts with the Wnt target genes and mammary bud markers *Dkk4* and *Fgf20* (Bazzi et al., 2007; Elo et al., 2017; Fliniaux et al., 2008), which, based on their expression in the Wnt-low milieu of *Eda*^*-/-*^ primary hair placodes (Fliniaux et al., 2008; Huh et al., 2013), respond to low levels of Wnt activation. This suggests that the mammary bud displays lower level of Wnt signaling activity than the hair placode or the enamel knot, possibly contributing to the different skin appendage identities. In line with this hypothesis, forced stabilization of β-catenin in the mammary bud led to ectopic expression of *Ascl4* along with several other hair placode markers including *Shh*, as well as *Fgf4* (Satta et al., 2023), an enamel knot –specific Wnt target gene (Jernvall et al., 1994; Kratochwil et al., 2002).

The bHLH transcription factors, including the Ascl family, play a central role in cell proliferation, cell fate specification, and differentiation (Ledent and Vervoort, 2001). This, together with the *Ascl4* expression pattern data, prompted us to analyze the phenotype of *Ascl4* null mice. However, we found no evidence for defective hair follicle morphogenesis, or tooth or mammary gland abnormalities. One possibility is that other Ascl transcription factors compensate for Ascl4. In developing teeth, our scRNA-seq data revealed co-expression of *Ascl5* with *Ascl4* in the enamel knot, a finding in line with Ascl5 (also known as AmeloD) immunostaining data (Chiba et al., 2019; He et al., 2019). Deficiency in *Ascl5* leads to enamel hypoplasia, yet the molar cusp pattern appears relatively normal (Chiba et al., 2019; Jia et al., 2022). We propose this could be due to redundancy with *Ascl4*. In the hair placodes, *Ascl5* and *Ascl2* were detected in the hair placodes by RNA-seq. Although both were expressed at low levels, we cannot exclude the possibility that they together, or alone, could function redundantly with *Ascl4*. An alternative option is that some other bHLH transcription factor compensates for loss of Ascl4. Future studies involving compound mouse mutants are warranted to address these possibilities.

## Materials & Methods

### Mouse strains

For *in situ* hybridization experiments, NMRI mice (Envigo) were used, unless stated otherwise. C57BL/6NJ-Ascl4^*em1(IMPC)J*^/Mmjax mouse line (MMRRC_#046169-JAX, here referred to as *Ascl4* KO), were obtained from the Jackson Laboratory and maintained on C57Bl/6J background. Transgenic mouse lines used in this study have been described elsewhere as indicated: *Eda* null mice (Pispa et al., 1999), *Ctnnb1*^flox3/flox3^ (Harada et al., 1999) and K14^Cre/wt^ mice carrying a knock-in of Cre in the *Krt14* locus (Huelsken et al., 2001). Embryos were staged according to limb development and other morphological criteria; noon of the vaginal plug was considered as embryonic day (E) 0.5. All mouse studies were approved and carried out in accordance with the guidelines of the Finnish National Experiment Board.

### Whole mount *in situ* hybridization

For whole-mount *in situ* hybridization (WMISH), embryos were fixed in 4% paraformaldehyde in PBS overnight, and dehydrated using a series of methanol. WMISH was performed manually as previously described (Mustonen et al., 2004). BM Purple AP Substrate precipitating Solution (Roche, Mannheim, Germany) was used to visualize the digoxigenin-labelled riboprobes. The following antisense RNA probes were used: *Ascl4* (Sulic et al., 2023), *Shh* (Echelard et al., 1993), *Dkk4* (Fliniaux et al., 2008), and *Wnt10b* (Wang & Shackleford, 1996). The samples were imaged using Zeiss 456 AxioZoom.V16 stereomicroscope with PlanZ 1.0x/C objective and Axiocam 305 color camera (Zeiss, Oberkochen, Germany), and handled using ZEN 2.3 lite software (Zeiss, Oberkochen, Germany).

### Hanging drop culture, and qRT-PCR

The hanging drop culture was performed as previously described (Pummila et al., 2007; Fliniaux et al., 2008). Briefly, E14.5 skins were dissected from NMRI embryos, and cut in half along the dorsal midline: one half was used as a control and the other one treated with CHIR99021. Samples were submerged in DMEM, 10% FBS, 1% penicillin-streptomycin supplemented with either 5 μM CHIR99021 (CT99021) (Sigma-Aldrich, St. Louis, MO) or the respective volume of DMSO as a vehicle control, and placed individually in 50μl hanging drops of under the lid of a 35 mm culture dish containing medium/PBS to prevent evaporation and maintained in a cell culture incubator at 37°C, 5% CO_2_ for 4 hours. Skin halves were collected directly to TRIzol reagent (Sigma-Aldrich, St. Louis, MO) and stored at -80°C until RNA extraction. Total RNA was extracted using the Direct-zol RNA MicroPrep kit (Zymo Research, Irvine, CA) according to manufacturer’s instructions. cDNA synthesis for qRT-PCR was done with iScript gDNA Clear cDNA Synthesis kit (Bio-Rad, Hercules, CA) according to the manufacturer’s instructions, using 1 μg of RNA as input, and diluted to 10ng/μl.

qRT-PCR was carried out with CFX96TM Real-Time System C1000Touch Thermal Cycler (Bio-Rad, Hercules, CA) using FAST SYBR Green master mix (Thermo Fisher), in triplicate wells. The following cycling protocol was used: Initial denaturation 3 min at 95°C, followed by 45 cycles of denaturation 10 s at 95°C and annealing/extension 50 s at 60°C. qRT-PCR data was analyzed with CFX Manager (Bio-Rad, Hercules, CA) software and fold changes were calculated using the ΔΔCt method (Livak & Schmittgen, 2001). These were normalized to *Hprt* gene. qRT-PCR primers were designed as following using NCBI Primer BLAST software: *Ascl4* (NM_001163614.2) (Forward primer: TGCTGACCCAAGGAGTGAAT; Reverse primer: CGCCTCATCAACGCTTGTAT), *Hprt* (NM_013556.2) (Forward primer: CAGTCCCAGCGTCGTGATTA; Reverse primer: TCGAGCAAGTCTTTCAGTCCT).

### Histology

For histology, tissues were fixed overnight in freshly prepared 4% paraformaldehyde in phosphate-buffered saline and embedded in paraffin. Transversal sections of 5μm were stained by hematoxylin and eosin using standard protocols and imaged using Axio Imager M.2 widefield microscope equipped with Plan-Neofluar 20x/0.5 objective and AxioCam HRc camera (Zeiss, Oberkochen, Germany) using bright field microscopy.

### Micro-Computed Tomography

Head samples were collected from 4-week-old wild-type and *Ascl4* KO mice and stored in 70% EtOH. To scan the teeth, heads were immobilized in 1% agarose. X-ray scanning was performed using Bruker 1272 μCT scanner at the μCT laboratory in the Department of Physics, University of Helsinki. The source voltage of 70 kV and source current of 142 μA was used with 0.5-mm aluminum filter. A voxel size of 4.5 μm and 18 μm for molars and incisors were used, respectively. For the reconstruction of the μCT scans, NRecon software (1.7.4.2) with the ring artefact correction of 18 was used. Tooth region was cropped and reconstructed as 8-bit tiff files in Fiji-ImageJ (version 1.53c), and segmentation was done using Avizo (release 9.0.1). For molars, the files were saved as .stl files in ImageJ (Fiji), and final 3D images were obtained using MeshLab (version 2021.10).

### Carmine Alum Staining

4-week-old (29 days) female mice were sacrificed, and the entire mammary gland 4 with fat pad was dissected out and spread on glass slides, fixed overnight in acid alcohol (2:8 mixture of glacial acetic acid and 100% ethanol), rehydrated through ethanol series, and stained in carmine alum (Sigma-Aldrich) overnight. Stained tissues were dehydrated in ethanol series, cleared in xylene, and mounted with Depex (BDH). Images were taken with Zeiss 456 AxioZoom.V16 stereomicroscope with PlanZ 1.0x/C objective and Axiocam 305 color camera (Zeiss, Oberkochen, Germany). Number of tips were quantified manually using Image J (Fiji).

### scRNA-seq data analysis

E14.5 hair placode-enriched (GSE212674) and E14.5 molar teeth (GSE142201) scRNA-seq datasets were analyzed as previously described (Hallikas et al., 2021; Sulic et al., 2023). Molar data was visualized using t-distributed stochastic neighbor embedding (tSNE) dimension reduction in R package Seurat v4.2.0.1. (Hao et al., 2021), R v4.3.1/RStudio v2023.09.1+494 (Posit Software, PBC). Molar epithelial cells were identified based on expression of *Krt14* and *Epcam*, subsetted and subclustered following Seurat’s documentation, using 40 dimensions and resolution of 0.2. Stacked violin plots were produced using scCustomize R package v1.1.3 (Marsh SE, 2021).

### Statistical analysis

For qRT-PCR data, quantification of mammary gland tips, and weight data, statistical analyses were performed using the GraphPad Prism 7 software and two tailed unpaired Student’s T test. p < 0.05 level of confidence was accepted for statistical significance. Accordance to Mendelian genetics was assessed using the GraphPad Prism 7 software and the Chi-square test, with p value <0.05 considered statistically significant.

## Acknowledgments

We thank Ms. Raija Savolainen for excellent technical assistance, Mr. Otto Mäkelä for support in realizing histology and *in situ* experiments, and past and present members of the Mikkola and Jernvall labs for stimulating discussions. Mouse studies were carried out with the support of HiLIFE Laboratory Animal Center Core Facility, University of Helsinki. This work was financially supported by the Sigrid Jusélius Foundation (MLM), the Finnish Cultural Foundation (VP), Ella and Georg Ehrnrooth Foundation (JPS), and the doctoral program in Oral Sciences of the University of Helsinki (AKH). The funders had no role in study design, data collection and analysis, decision to publish, or preparation of the manuscript.

---

**Supplementary Figure 1.**
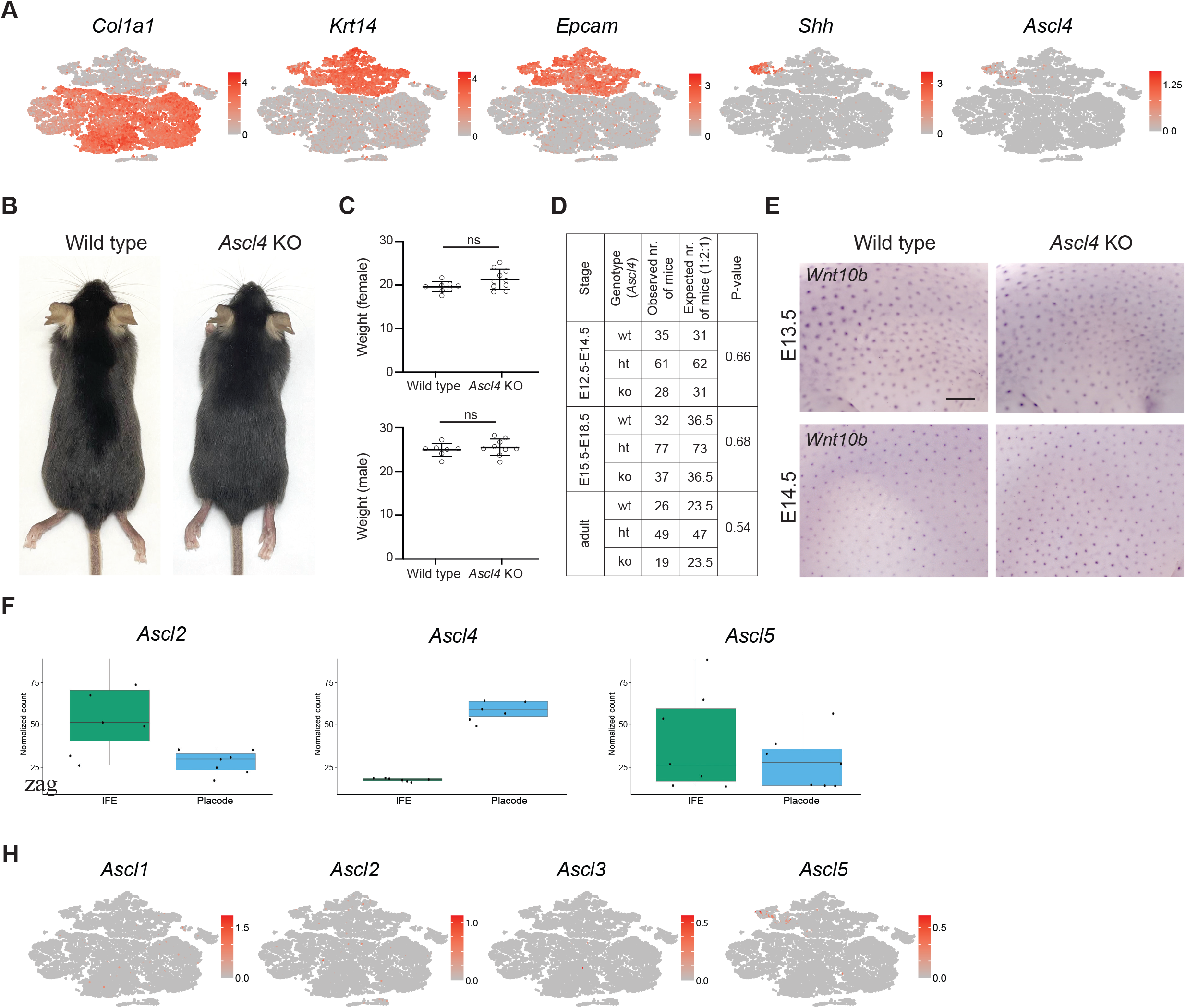
Expression of *Ascl4* and phenotype analysis of *Ascl4* mice. **(A)** tSNE plots showing expression of the mesenchymal marker *Col1a1*, the epithelial markers *Krt14* and *Epcam*, and the enamel knot markers *Shh*, as well as *Ascl4* in E14.5 molar tooth scRNA-seq dataset from Hallikas et al., 2021. Colors span a gradient from red (high expression) to grey (low expression) for each gene. **(B)** Representative images of wild-type and *Ascl4* KO mice (n=8 each) at 8 weeks of age. **(C)** The body weight of 8-week-old male and female *Ascl4* KO (n_male_=9; n_female_=10) and wild-type mice (n_male_=7; n_female_=8). Data are shown as mean ± s.d. Statistical significance was assessed with two-tailed unpaired Student’s t-test. ns, not significant. **(D)** The number of mice of various genotypes obtained from the *Ascl4* heterozygous matings is shown. The expected number was calculated according to the total number of mice born and based on the Mendelian 1:2:1 ratio. E12.5-E14.5 and E15.5-E18.5 embryos were pooled for statistical analysis. Statistical significance was assessed with Chi-square test. wt, *Ascl4*^+/+^; ht, *Ascl4*^+/-^, ko, *Ascl4*^-/-^. **(E)** Expression of *Wnt10b* was analyzed by whole mount *in situ* hybridization in E13.5 (top panel) and E14.5 (bottom panel) embryos (n=4 each). Scale bar, 500 μm. (F) Box plots for *Ascl2, Ascl4* and *Ascl5* showing normalized count distributions between interfollicular epithelium and placode samples in E14.5 hair placode bulk RNA-seq dataset (7 biological replicates) from Sulic et al., 2023. IFE, interfollicular epithelium. **(H)** tSNE plots showing expression of the transcription factors *Ascl1, Ascl2, Ascl3* and *Ascl5* in E14.5 molar tooth scRNA-seq dataset from Hallikas et al., 2021. Colors span a gradient from red (high expression) to grey (low expression) for each gene.

